# Advanced coarse-grained model for fast simulation of nascent polypeptide chain dynamics within the ribosome

**DOI:** 10.1101/2025.04.08.647858

**Authors:** Shiqi Yu, Artem Kushner, Ella Teasell, Wenjun Zhao, Simcha Srebnik, Khanh Dao Duc

## Abstract

The nascent polypeptide exit tunnel (NPET) is a sub-compartment of the ribosome that constrains the dynamics of nascent polypeptide chains during protein translation. Simulating these dynamics has been limited due by the spatial scale of the ribosome and the temporal scale of elongation. Here, we present an automated pipeline to extract the geometry of the NPET and the ribosome surface at high resolution from any ribosome structure. We further convert this into a coarse-grained (CG) bead model that can be used in molecular simulations. This CG model more accurately captures NPET geometry than previous representations and allows for the simulation of co- and post-translational processes that are computationally prohibitive with all-atom approaches. In particular, we illustrate how the CG model may be used to simulate the elongation dynamics of the nascent polypeptide and its escape post-translation, as well as evaluate free energy landscapes and examine the influence of electrostatics on the nascent polypeptide escape.

**SIGNIFICANCE:** The translation of nascent polypeptide chains is mediated by the ribosome, with interactions between the protein and the nascent polypeptide exit tunnel (NPET) impacting the process. However, modeling and simulating protein elongation and its escape from the NPET remain challenging due to computational limitations in spatial and temporal resolution. Here, we develop a computational pipeline for generating coarse-grained (CG) models of the NPET and ribosome surface from any ribosome structure that allow for both effective and accurate computer simulations. We demonstrate how the model can be implemented for various simulations, including the elongation dynamics of the nascent polypeptide and escape of the chain post-translation, as well as in estimating free energy landscapes and examining the impact of the charged environment on the escape time.

## INTRODUCTION

The ribosome is a large RNA-protein complex that mediates protein synthesis from messenger RNA (mRNA) during translation. This process requires the formation of a peptide bond at the peptidyl-transferase center (PTC), which is deeply embedded within the large ribosomal subunit (1). As peptides get added, the nascent polypeptide chain passes through the nascent polypeptide exit tunnel (NPET). The NPET is a narrow conduit, approximately 100 Å length, that connects the PTC to the ribosome’s solvent-exposed surface (2). Over the past decades, experimental efforts have focused on quantifying interactions between the nascent chain and ribosome and elucidating their impact on protein folding and translation arrest (3–6) to understand how NPET dynamically influences protein synthesis (7, 8). Given the tunnel’s dimensions and the complexity of these interactions, molecular dynamics simulations have emerged as both a complement and alternative to experimental methods, providing atomic-level insights into nascent chain interactions at the NPET (9–12).

Simulations involving internal and external features of ribosomes are challenging due to the sheer size a complexity of the ribosome. Bacterial and eukaryotic ribosomes measuring about 20 nm and 30 nm, respectively (13, 14). As such, running even a 10-nanosecond ribosome simulation on a single GPU may take a full day, and microsecond-scale simulations may extend over several months (9). Whole-ribosome simulations therefore remain exceedingly rare (9) and most molecular dynamics (MD) studies focus exclusively on the tunnel region (15–20). Yet, even these reduced models comprise tens of thousands of atoms (20) and create a substantial computational burden. Several other studies employ cylindrical approximations with varying radii to model the exit tunnel and examine protein escape dynamics (21–23). However, such studies risk oversimplifying some potentially important geometric features of the NPET (24). Moreover, most models fail to consider the ribosome’s solvent-exposed surface, which is known to modulate folding and translation dynamics of the nascent polypeptides as they emerge from the NPET (4, 25, 26). Notably, intermolecular interactions between the ribosome and specific amino acid (aa) residues at the N-terminus are known to perturb co-translation folding (27) and regulate translation fidelity (28) of nascent chains, and hence should not be neglected in models aiming to capture the ribosome’s role in modulating translation dynamics. All-atom MD simulations have historically been limited to treating one or few species due to their computational expense.

However, the recent surge in high-resolution ribosome structures obtained through cryo-electron microscopy (29) has revealed unprecedented heterogeneity across species and domains of life (30). Comparative studies have demonstrated how some important structural variations in the NPET of bacteria and eukaryotes significantly impact nascent chain dynamics (19, 24). This structural diversity amplifies the need of alleviating computational costs and facilitating efficient modeling of the NPET environment. This expanding structural dataset can be leveraged to comprehensively explore the functional role of the NPET across evolutionary lineages.

To address these computational limitations, we developed a pipeline to construct coarse-grained (CG) models of NPET and ribosome surface from any given ribosome structure that simplifies and reduces the complexity of simulating nascent polypeptide chain dynamics. In particular, when utilized with coarse-grained models of proteins (31), the model allows for simulating dynamics at the time scale of protein translation. Our approach converts empty space voxels from the ribosome 3D structure into a 3D mesh of the NPET, that is subsequently used to create an efficient CG model for studying translation-related processes. We benchmark our model against established tunnel representations, evaluating its geometric accuracy and computational efficiency in simulating nascent polypeptide dynamics. We then demonstrate the versatility of the model through various applications, including simulating co-translational and post-translational processes, as well as comparison of energy profiles, and evaluation of escape kinetics of charged polypeptide chains, showing its potential as a tool for future investigations of nascent polypeptide chain interactions and dynamics across diverse ribosome structures.

## METHODS

### Modeling the NPET interior and ribosome exterior surfaces

To build a CG model of the NPET and ribosome, we first extract the surface of the ribosome and the interior surface of the NPET as regular meshes. For any arbitrary atomic model of the ribosome (obtained from the PDB or other database (29) as a .`cif` file), we run the following protocols, illustrated in Figure S1:

### Exterior ribosome surface

Briefly, we obtain the atomic model of the ribosome deposited in the PDB and parse its (32) .`cif` file to extract 3D coordinates of atoms. This constitutes the initial pointcloud. A pointset is extracted from that cloud that represents only the surface atoms via Delaunay triangulation (33). Further, orientation vectors known as normals are estimated (34) on this surface pointset to establish what constitutes *the interior* and *the exterior* in space relative to the surface pointset. Poisson surface reconstruction is applied to the normal-estimated pointset which yields a smooth mesh of the ribosome surface. Examples of corresponding structures generated during the stages of the modeling of the exterior ribosome surface are shown in Figure S1a.

### Interior NPET surface

To model the NPET, we run the following six steps (with visualization shown in Figure S1b.

1. The locations of the PTC and the *Constriction Site* are retrieved from the API of the riboxyz project: api.ribosome.xyz (for a precise definition of how their coordinates are calculated as well as, see SI file)
2. The pointcloud of the given ribosome structure is obtained from the corresponding PDB file as described in the previous section (*Exterior ribosome surface*). Any fragments that might be occupying the tunnel (*e*.*g*. nascent peptide chains and ligands) are removed as needed.
3. Using the PTC and the constriction site coordinates as the main axis, we construct a cylindrical mask with radius of 35 Å and height of 120 Å with the PTC at the base. These dimensions are heuristic but conservative enough to contain any ribosome tunnel across the tree of life. This cylindrical mask is mounted on the axis defined by the PTC and the *Constriction Site* and applied to the ribosome pointcloud.
4. Atoms contained in the mask are projected onto a binary voxel grid of the same dimensions as the cylindrical mask. The positions of atoms which belong to the mask are used to activate (set value to 1) voxels whose centers fall within a sphere of radius 2 Å from the given atom positions. This 2Å sphere is a simplistic representation of the van der Waals radius of an idealized atom. The voxel grid is clipped with the surface of the whole ribosome (produced above in *Exterior ribosome surface*) to discard the inactive voxels (representing empty space past the tunnel exit port).
5. At this point the cylindrical voxel grid contains only values 0 and 1, where inactive (0-value) voxels correspond to a mixture of the NPET interior space and inter-polymeric space pockets and active (1-value) voxels represent space occupied by atoms. We use DBSCAN (36) density-based clustering algorithm to distinguish the central and biggest cluster corresponding to the NPET interior from the other pockets. We run DBSCAN two times, first on the entire grid (with default parameters *ε* = 5.5 Å and *min*_*san ples* = 600) to detect major clusters, and second on the central cluster (*ε* = 3 Å and *min*_*san ples* = 120) to sharpen the cluster revealing atomic-scale features. For a structural interpretation of DBSCAN parameters, see SI File.
6. As in the previous section, we estimate the normals on the surface of the NPET interior voxel cloud. Poisson Surface reconstruction (35) is likewise applied to this final voxelcloud to produce a smooth watertight mesh.

### Coarse-grained representation of the ribosome tunnel and surface

The aforementioned mesh structure serves as the initial scaffold for the CG model, with CG beads placed at the centroid of the triangular mesh. The beads representing the NPET are placed along the surface normal at a distance corresponding to the radius of the bead to properly capture the cavity. For the ribosome surface, the CG beads are placed directly on the mesh. If necessary, additional CG beads can be added or removed on the mesh vertices or edges. The centroids and normal vectors of the triangle meshes are computed using the PyMeshLab library in Python (37). LAMMPS “delete_atoms” command (38) is used to delete atoms to ensure an open NPET exit port and reduce the computational costs by deleting overlapping CG beads.

### Mapping effective charges onto the CG beads

The effective potential at the center of each CG bead at the NPET, *j*, is calculated using the screened potential, *ϕ* _*j*_, summed over all charged residues (lysine, arginine, glutamate, and aspartate) and nucleic acids within a certain distance (*e*.*g*., 50 Å) of the tunnel centerline, as

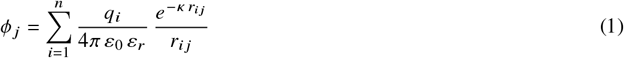

where *r*_*ij*_ is the distance between the charged atom, *i*, and the CG bead, *j*. *q*_*i*_ is the atom charge, *ε*_0_ is the vacuum permittivity, *ε*_*r*_ is the dielectric constant, and *κ* is the inverse Debye length. The coordinates of the charged residues are extracted using PyMOL and, for simplicity, assigned to their respective *C*_*α*_ atoms (39). The positions of negatively charged rRNA atoms are placed on the backbone phosphorus atoms. The value of *κ* is taken to be 0.14 Å^−1^. The dielectric constant, *ε*_*r*_, can be set to unity for explicit solvent simulations, can be used to best fit the experimental data (3), or can be estimated in the range of 4 to 80 (40). The effective charge of each CG bead of NPET is then evaluated from 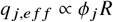, where *R* is the bead radius, set as 1.8 Å (41). The proportionality constant is chosen to align the induced potential along the tunnel centerline of the CG model with that from the atomistic model using least squares optimization.

While the effective charge of each CG bead of the ribosome surface can be similarly calculated as above, we alternatively propose to calculate the effective electrostatic potential for implicit solvent simulations using Adaptive Poisson-Boltzmann Solver (APBS) in PyMOL (42, 43). This approach accounts for the distinct solvent environment outside the ribosome compared to the tunnel interior. The calculations of the linearized Poisson–Boltzmann equation (PBE) (43) are performed at 0.15 M ionic strength in monovalent salt, 310K, solute dielectric constant of 4 and a solvent dielectric constant of 80 (44). The grid spacing is set to 1 Å. All other parameters of the APBS input file are kept at the default settings. The effective potentials at the centers of the CG beads are then obtained through three-dimensional cubic interpolation of the grid-based data from APBS.

### Dynamic simulations of nascent chains

A simplified CG protein model is used in this study to simulate nascent chain dynamics to illustrate the CG tunnel and ribosome model. Each residue is modeled by a triplet of beads representing the backbone units, where the central bead of each triplet is designated as either neutral, positively charged, or negatively charged. Lennard-Jones (LJ) potential is used to model the short-range intermolecular interactions between non-bonded residues (45). Intramolecular interactions are modeled with harmonic bond and angle potentials, and the dihedral force between neighboring residues. The screened Coulomb potential is used for electrostatic interactions.

To simulate post-translational dynamics within the NPET, CG beads of the nascent chain are initially aligned along the tunnel centerline with the C-terminus positioned at the PTC. The attached polypeptide chain is first equilibrated as the average internal energy remains constant for 50,000 timesteps. The C-terminus is then released. Co-translational dynamics are modeled by simulating the intermittent addition of beads at the PTC at a constant rate. The chain is advanced by a distance equal to half the bond length towards the exit to introduce new beads. During each interval between bead additions, the nascent chain is attached at the PTC. This process is repeated iteratively until the desired chain length is reached. For details on the MD protocol and force field parameters, see SI file.

### Datasets

The structures used to construct the mesh of the NPET and ribosome surface in this paper are high-resolution atomic models of the ribosome from *H*.*sapiens* and *E*.*coli*, deposited in the Protein Data Bank (PDB) with codenames 4UG0 and 6WD4 respectively.

### Software and implementation

Our pipeline runs in Python, using a combination of packages including numpy (46), pyvista (33), scipy (47), with the code and illustrative notebooks publicly accessible on GitHub. MD simulations were conducted using LAMMPS (released version: 29 Aug 2024) (48) on a single CPU compute node with a consumer-line PC with 1.4 GHz Quad-Core Intel Core i5 and 8GB RAM memory. The coordinate extractions of the tunnel centerline, charged residues and atoms were performed in PyMOL (49). Visualizations of the coarse-grained models and nascent chain dynamics are carried out with VMD (50), and OVITO (51). Cross-sectional views of the ribosome models are generated using ChimeraX (52). All downstream analyses were conducted using custom Python scripts.

## RESULTS

### Coarse grained modeling of the ribosome surface and NPET

We developed a computational pipeline to generate coarse-grained models of the NPET and ribosome surface from any given structure, as shown in Figure 1a and detailed in the Methods section for a more detailed visualization, see also Supplementary Figure S1. Beginning with an atomistic model of the ribosome, the exit tunnel is first identified based on landmarks that define the PTC and constriction site. Atoms within a cylindrical volume surrounding the tunnel are used to extract the tunnel surface as a mesh through DBSCAN clustering and Poisson surface reconstruction (Figure 1b). CG beads are then placed on the mesh to form a CG representation of the tunnel surface. The ribosome surface is similarly captured (Figure 1c). The CG assembly, integrating both the tunnel and ribosome surface can be seen in 3D in Supplementary Video S1.

**Figure 1.**
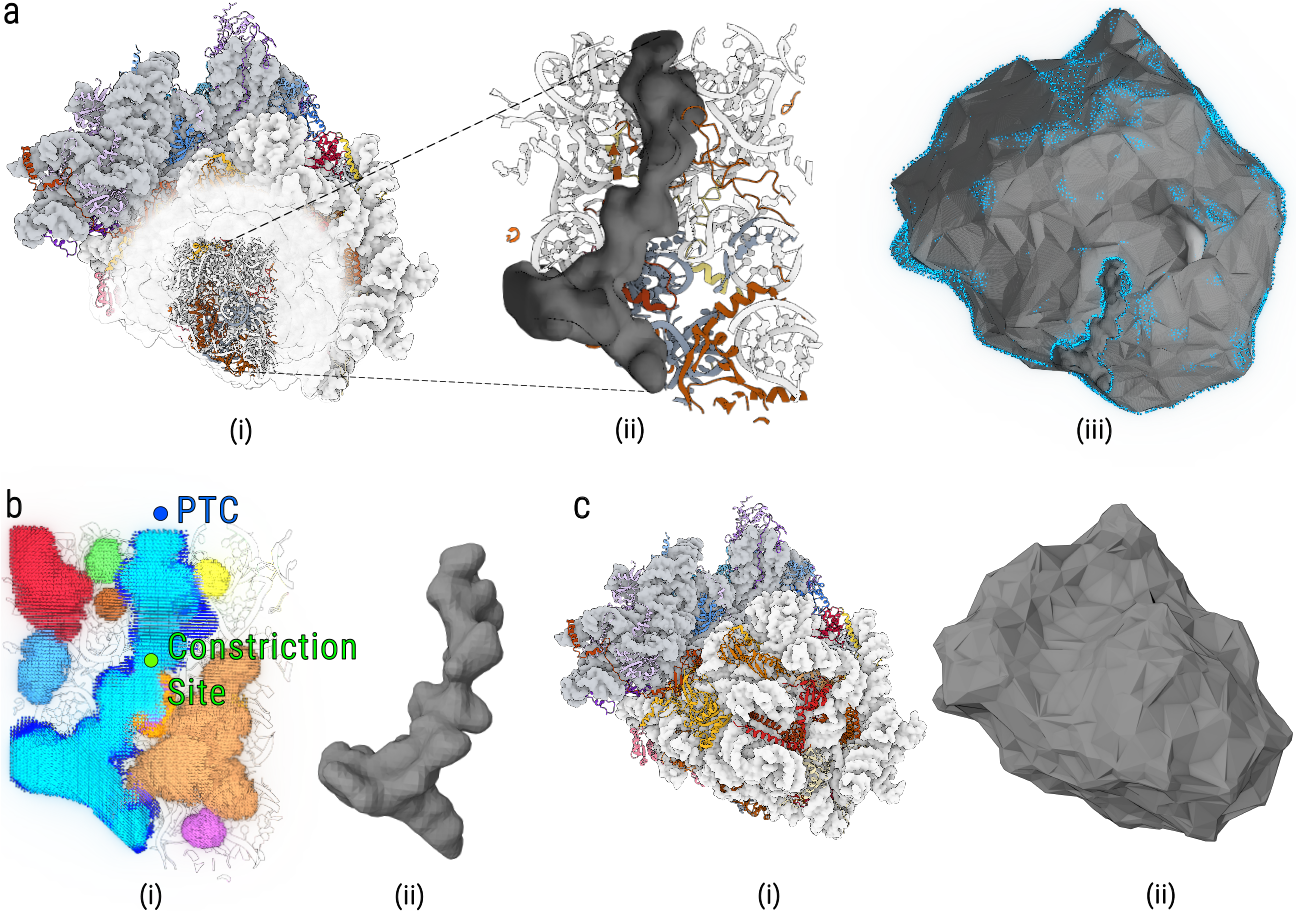
Construction of CG simulation environment illustrated with ribosome structure 4UG0. **(a)** General illustration showing the *(i)* ribosome structure, *(ii)* NPET and *(iii)* ribosome surface. For a better visualization of the tunnel position and geometry, only selected representative beads of the CG model are shown as blue spheres in *(iii)*. **(b)** After *(i)* clustering empty voxels, we *(ii)* build the tunnel mesh. **(c)** Mesh produced from the *(i)* ribosome structure using *(ii)* Poisson surface reconstruction.

### Modeling effective charges

As an RNA-protein complex enriched in charged residues, the ribosome will have significant electrostatic interactions with charged polypeptide chains (16, 53). To include these interactions in our CG model, we mapped the effective charges for both the ribosome tunnel and surface onto the CG beads. For the beads at the NPET, we used the ribosomal charged residues and nucleic acids within a certain distance of the NPET, and calculated the associated screened potential at each bead (see Eq. (1)), as described in the Methods section. For the ribosome surface, we used the APBS solver in Pymol (see Methods section) to account for the solvent in our calculation of the effective potential. Figure 2a illustrates the charge distribution of the CG model for the 4UG0 ribosome and NPET (see Figure S2a for 6WD4), with both showing predominantly negative charge. Scattered positively charged regions on the ribosome surface are primarily associated with ribosomal proteins (43) as has been observed in cryo-EM ribosome structures.

**Figure 2.**
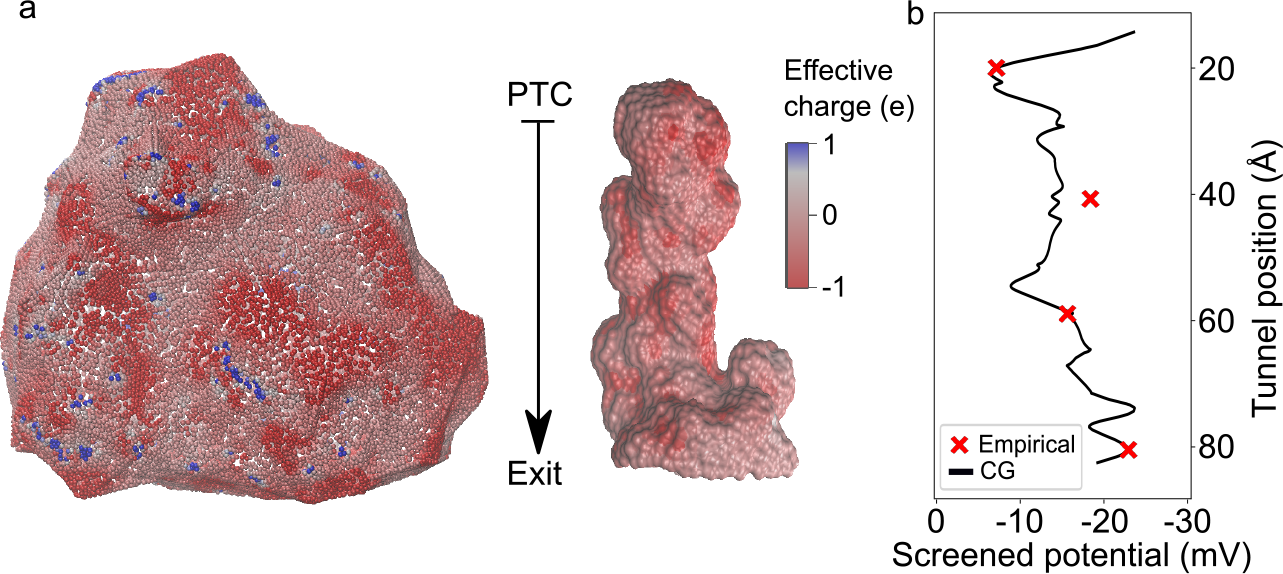
Coarse-grained models of the ribosome surface and tunnel with mapping of effective charges. **(a)** Visualization of effective charges mapped onto the ribosome surface and the exit tunnel of 4UG0. Blue and red beads represent positively and negatively charged beads, respectively. The color scale range is adjusted to improve visualization clarity. **(b)** Electrostatic potential along the tunnel centerline in the archaeal ribosome 1JJ2 derived from the CG model (black curve) compared with experimental data (3) (red crosses).

To evaluate the accuracy of our mapping, we compared the electrostatic potential profiles along the tunnel centerline generated by our CG model with experimental measurement measured at specific positions of the tunnel for an archaeal ribosome (3) (Figure 2b), as well as the electrostatic profile obtained from atomistic simulation of the bacterial ribosome (4V5D) (44) (Figure S2b). The results indicate that the profile from the CG model aligns closely with empirical measurements (see Figure 2b), and shows the same trend as for the atom-based model. We also noted some more significant deviation between the CG and atom-based profiles at the exit port (Figure S2b), that can be attributed to the inclusion of magnesium ions and TIP3P waters in the atom-based model (44).

### Comparison with other models

The extent to which the CG model more accurately captures the NPET geometry was evaluated by comparing it to the model obtained by running MOLE, a cavity search algorithm (54, 55) that was used in previous recent geometric studies of the NPET (24, 56). As this algorithm yields a set of spheres lining along a centerline from the PTC to the exit port, it likely fails to capture irregularities along cross sections of the NPET. Taking ribosomes 4UG0 (*H*.*sapiens*) and 6WD4 (*E*.*coli*) as examples from different life domains, we used MOLE2.5 (54, 55) to extract the NPET model and compared the volume produced with our method. To do so, we also divided the tunnel into three regions shown in Figure 3a, namely the upper, lower and exit regions. These regions are separated by the constriction site (see Methods), and conserved residues K68 in uL23 (bacteria), or R28 in eL39 (eukaryotes) (24, 57). As shown in Figure 3a for 4UG0, the CG model encompasses a volume that is significantly larger, with 3.2 ×10^4^ Å^3^ (2.9 ×10^4^ Å^3^ for 6WD4, see Figure S3a), compared to 7.3× 10^3^ Å^3^ (9.5× 10^3^ Å^3^ for 6WD4) for the spherical model. While the upper and lower regions in the CG model are 53% and 82% larger (45% and 71% for 6WD4), respectively, we observed that most of the volume discrepancy concentrates in the exit region, where the difference yields an order of magnitude difference, with 1.6 × 10^4^ Å^3^ for the CG model compared to 1.5 × 10^3^ Å^3^ for the spherical (1.2 × 10^4^ Å^3^ and 2.7× 10^3^ Å^3^ for 6WD4). The smaller exit volume observed in the spherical model results from the probe-based algorithm used for extracting cavity terminating earlier, as it could otherwise generate artificial paths (54). Overall, the CG model provides a more comprehensive representation of the NPET’s internal space, particularly at the exit port and interface with the external surface. A visualization comparing the volumes of the two tunnel models is provided in the Supplementary Video S2.

**Figure 3:**
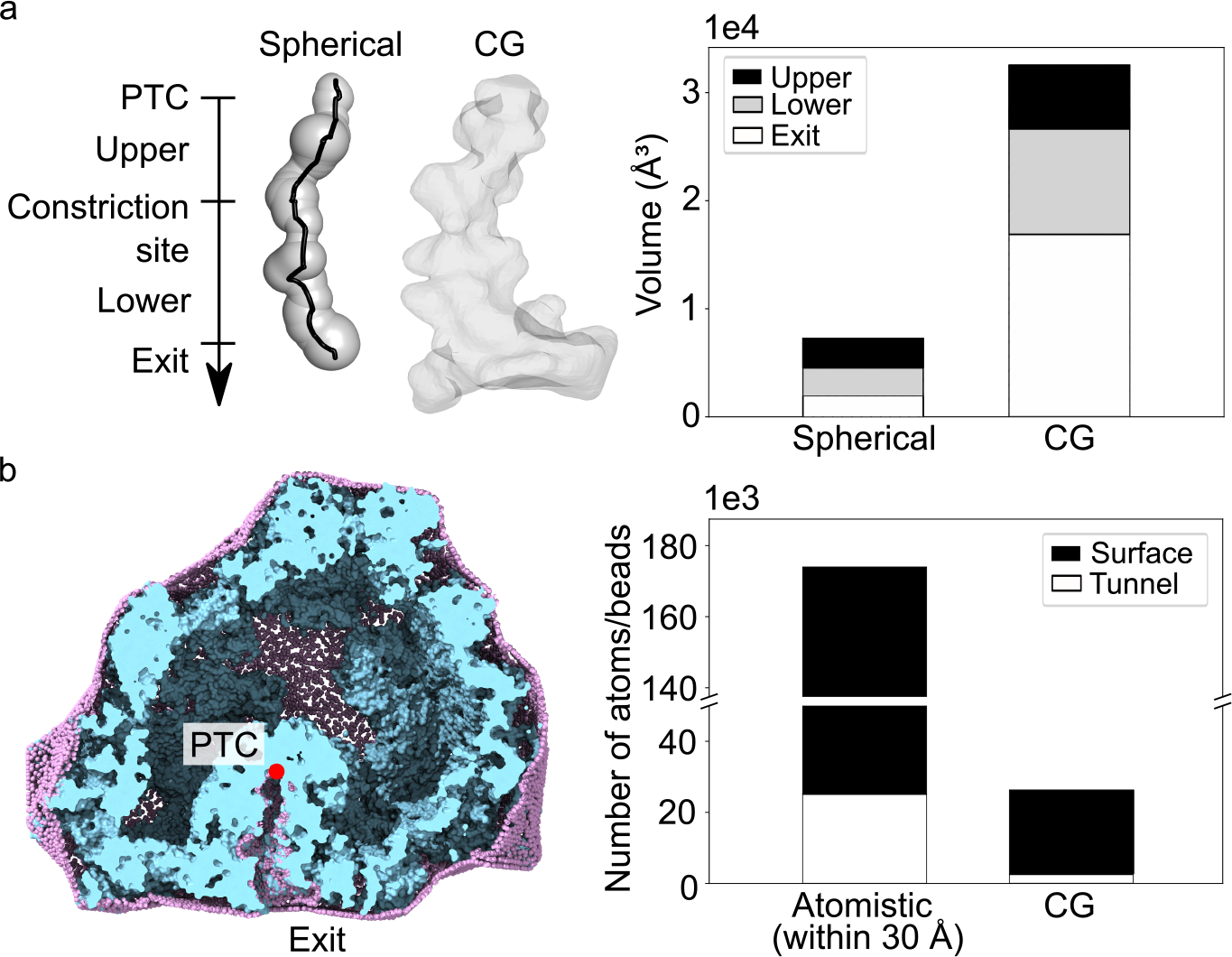
Comparisons of the CG model with the spherical and all-atom model for 4UG0. **(a)** The left panel shows the MOLE-generated spherical tunnel model (left) and the CG tunnel model in mesh (right). The tunnel is divided into three regions. The upper, lower and exit region are separated by the constriction site and conserved residues in uL23 or eL39. The right panel compares the volume of the spherical and CG tunnel models in the upper, lower and exit regions. **(b)** Perspective view of a cross-section of the ribosome, showing the atomistic model including all atoms within 30 Å of the ribosome surface (cyan), and the CG ribosome model (purple). Lighter shades of blue indicate atoms closer to the viewer, while darker shades represent atoms farther away. The PTC is highlighted as a red dot. The right panel shows the comparison of the number of beads or atoms in the tunnel (white) and ribosome surface (black) regions for both models. In the atomistic model, overlapping atoms between the two regions are excluded in the atom count for the surface region.

To evaluate the potential benefit of the CG model in MD simulations, we further quantified the number of atoms that is typically required to simulate the ribosome environment in an atomistic representation. For 4UG0 we extracted all atoms located within 30 Å of the ribosome surface and the tunnel centerline (see Figure 3b). In the NPET region, the number of atoms required for 4UG0 is approximately 25,000 atoms (24,000 atoms for 6WD4). While our CG tunnel model contains 2,500 beads, that were set with a radius of 1.8 Å to be watertight (2,300 beads for 6WD4). When considering an implicit solvent CG protein model like that used in our following simulations, this number can be further reduced to *∼* 1,600. For the ribosome surface region, our CG model includes approximately 24,000 beads – about 80.5% reduction compared to the atomistic representation, which contains *∼* 123,000 atoms (22,000 and 95,000 atoms for 6WD4, around 76.8% reduction, see Figure S3b). This substantial decrease in system size significantly enhances computational efficiency and makes the model well suited for studying long-timescale processes, such as nascent chain elongation.

### Simulations of nascent chain dynamics

In this section, we demonstrate two applications of our CG model for MD simulations using a simplified nascent chain model. The nascent chain is represented using a bead-spring model where each residue comprises a triplet of beads representing backbone units. To isolate geometric and entropic effects, simulations were performed in a neutral CG ribosome environment. Detailed parameters are provided in Methods and SI file. Dynamic trajectory visualizations are provided in Supplementary Videos S3 and S4.

We first simulated the co-translational dynamics (elongation) of the polypeptide in the CG model of 4UG0 encompassing both the NPET and ribosome surface. To do so, we added each residue to the growing chain at the PTC (see Methods section) at 1× 10^4^ time step intervals. Considering a typical elongation rate of ~ 10 codons per second measured in yeast (58), this corresponds to a time step of ∼ 10 *µs* – a timescale characteristic of molecular CG simulations (31). Figure 4a shows snapshots of the elongation simulations. The 20-mer chain is is seen to be contained within the NPET, which is consistent with previous findings that ribosomes can accommodate up to 50 residues internally (59, 60). As elongation progressed to 80 and 100 residues, we see the N-terminal portion interacting with the ribosomal surface before extending into the solvent environment where folding might initiate.

**Figure 4.**
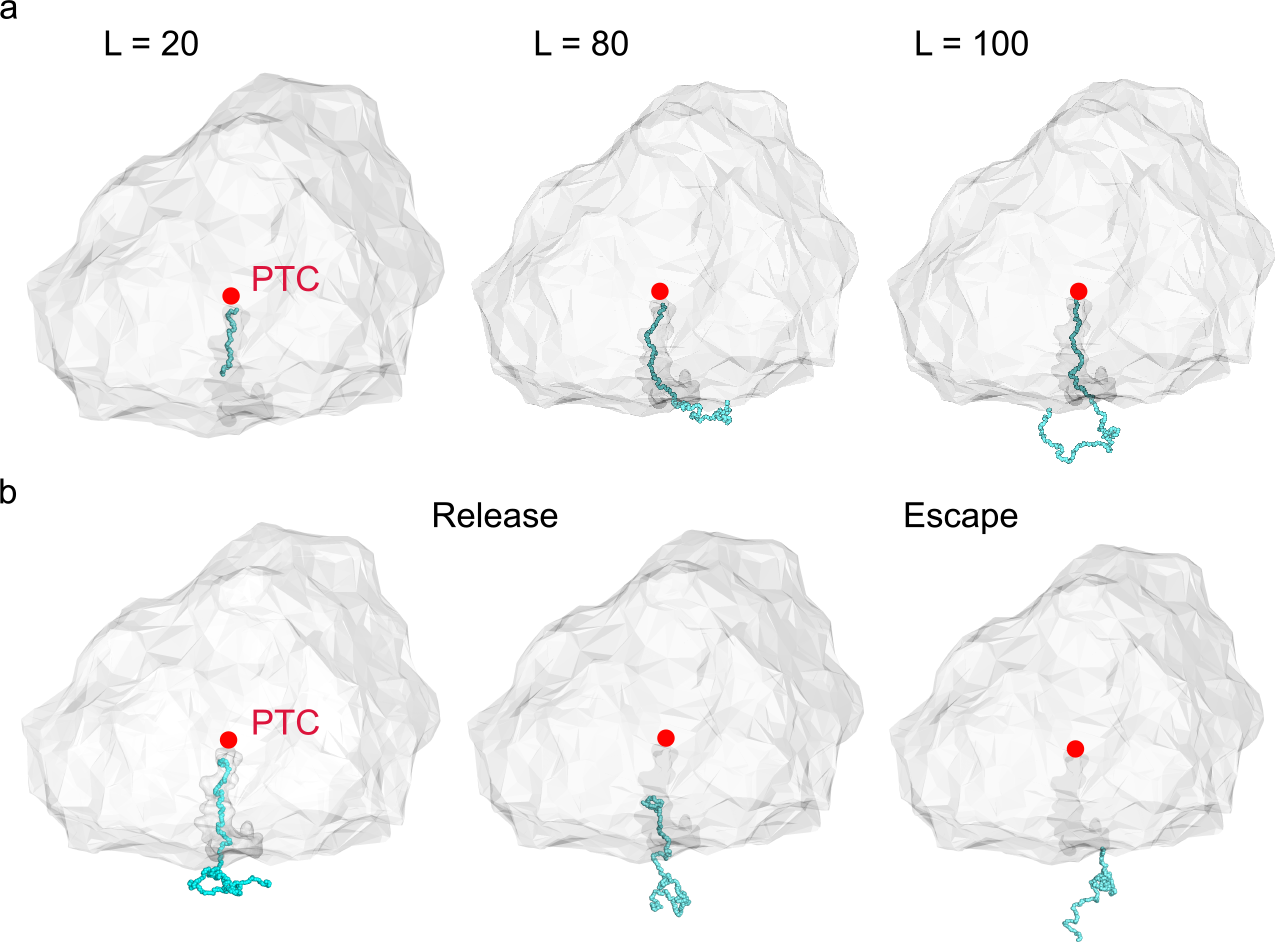
Dynamic simulations of polypeptides within the coarse-grained ribosome model of 4UG0. **(a)** Co-translational process of polypeptide elongating to 20, 80, and 100 amino acid residues. The ribosome surface is modeled to simulate interactions with the protein segment extending beyond the tunnel. Length *L* represents the number of amino acid residues elongated. **(b)** Post-translational dynamics of a 100-mer polypeptide, illustrating its release from the PTC to its full exit from the tunnel (left to right). The PTC is highlighted as a red dot.

Next, we simulated the post-translational escape process of polypeptides from the CG ribosome model of 4UG0, and Figure 4b illustrates the escape snapshots of a 100-mer polypeptide. The escape time of nascent chains can greatly vary within a 1000-fold range (16), with short polypeptides that fully accommodated within the tunnel before release exhibiting particularly extended times compared to other chain lengths (see next section for a more detailed explanation). For example, the simulation of the 20-mer polypeptide completed in approximately 25 minutes, from the equilibrated conformation of the peptide chain with its C-terminus attached to the PTC, until complete escape. While for the majority of the simulation, the chain remains confined within the upper tunnel region (see Figure S4 and next section), suggesting that the translocation of short polypeptide within the tunnel can be strongly hindered by the constriction sites (21). Table 1 summarizes the simulation time steps and corresponding CPU runtimes for both co- and post-translational processes. The simulation of a 500-mer chain elongation completed in approximately 4 hours, demonstrating that our CG model enables long-timescale protein translation simulations to complete within hours, even on a single CPU core.

**Table 1:** Simulation and CPU times for co- and post-translational processes for different lengths of polypeptides in the coarse-grained ribosome model of 4UG0.

| Simulations | Time step [A.U.] | CPU time [s] |
| --- | --- | --- |
| **Elongation**<br>(50-mer) | $1.6 \times 10^6$ | 1,206 |
| **Elongation**<br>(100-mer) | $3.2 \times 10^6$ | 2,282 |
| **Elongation**<br>(500-mer) | $1.6 \times 10^7$ | 15,040 |
| **Post-translation**<br>(tunnel escape) | 10,000 | 2.71 |

### Comparison of energy profiles

Focusing on the post-translation escape process illustrated in our second example, we study the nascent chain dynamics and free energy landscape the protein experiences in the NPET. Previously, we employed a simplified cylindrical model (21) to analyze nascent chain free energy profiles within the tunnel. While this cylindrical model captures the average geometric properties of the NPET across 20 eukaryotic species, significant variations can be observed across species (24), potentially leading to differences in energy profile. Like in our previous study, we considered here 12-mer and 20-mer polypeptides in the CG tunnel model of 4UG0, and calculated their free energy profiles along the tunnel axis. Free energy (*f*) landscapes for each polypeptide length were derived from simulations, by summing over the canonical partition function along the tunnel, with 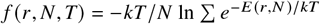 (see (21)), where the sum is over all configuration with C-terminus at position *r* along the tunnel, *k* is the Boltzmann constant, *N* is the number of beads in the polypeptide, and *E* is the energy of each microstate.

Figure 5a compares the energy profiles between the previous simplified cylindrical geometry and the current CG model. While both approaches reveal two distinct low-energy states, the energy wells in the CG model are notably shallower with lower energy barriers compared to the cylindrical representation. This difference highlights how a more accurate CG model provides more nuanced insights into the energetic landscape governing nascent chain transit through the ribosomal tunnel, with differences across species that could not be captured using a simplified geometry.

**Figure 5.**
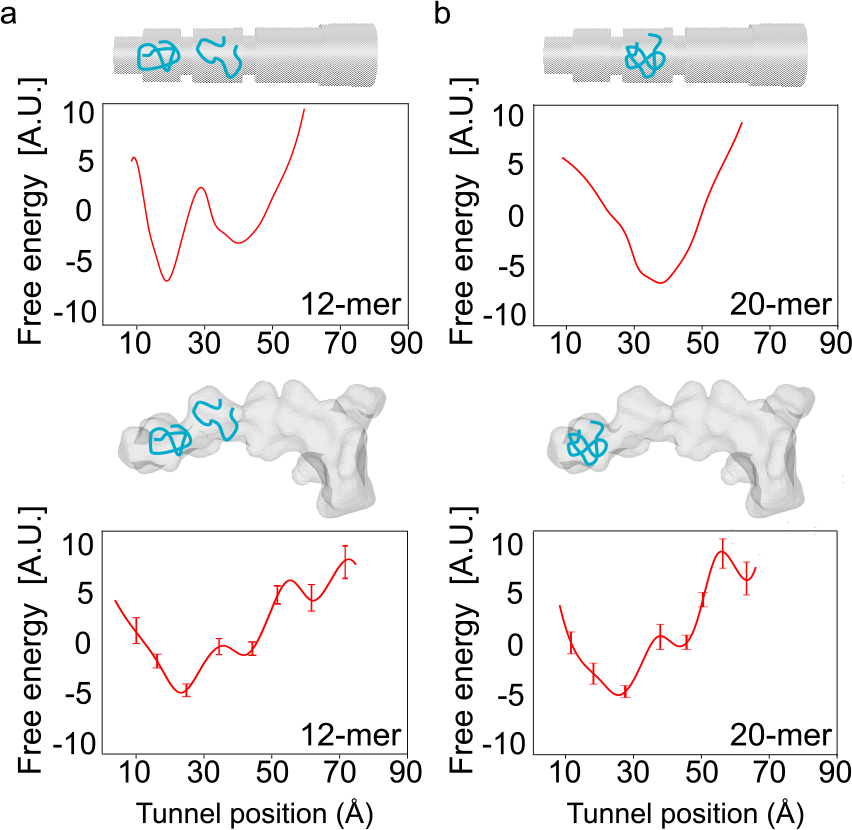
Comparison of free energy landscapes of **(a)** 12-mer and **(b)** 20-mer polypeptide in the cylindrical (upper) (21) and coarse-grained model (lower) of the neutral eukaryotic tunnel 4UG0. The energy is calculated from the sum over the partition function at each position along the tunnel using the C-terminus position. The error bars are calculated based on one hundred independent runs. A Savitzky-Golay filter is applied to reduce the noise.

For the 20-mer polypeptide (Figure 5b), we observed more significant qualitative differences between the two models. The cylindrical model only displayed a single energy well at the constriction site, whereas the CG model revealed a two-potential well profile again, with the deeper well located in the upper tunnel region. We attribute this discrepancy to volume underestimation in the cylindrical model’s upper tunnel region (as evidenced in Figure 3a), making it insufficient to accommodate the polymer chain. In contrast, the CG model demonstrates that the upper tunnel region can effectively trap both 12-mer and 20-mer polypeptides—a finding consistent with previous observations that short proteins of these lengths can initiate folding near the PTC (19, 61). While both models support the general conclusion that tunnel geometry impedes 20-mer polypeptide escape, it is interesting to note that their biophysical interpretations differ for 4UG0, highlighting again the importance of using a more detailed representation of the NPET.

### Escape of charged polypeptide

We finally illustrate how the CG ribosome model can be used to examine the effects of electrostatics on nascent chain escape dynamics, as previously done in *E*.*coli* (16). To do so, we constructed 80-mer polypeptides with 25 distinct charge patterns, each consisting of a different number of positively and negatively charged residues. For each non-neutral pattern, five sequences were generated by randomly distributing the charges within the C-terminal 30 residues, resulting in 120 unique charged sequences, along with one neutral sequence. Each sequence was simulated in 10 independent runs. Figure 6a shows the distribution of escape times, *T*_*e*_, from all simulations, normalized by the shortest escape time, *T*_*e,min*_. The distribution spans a wide range of escape times with a peak around 10, similar to the findings of Nissley *et al*. (16). We also note a long tail associated with slow escape events. In Figure 6b, we plot the normalized mean escape times for each charge pattern, revealing a clear charge-dependent bias, with the slow escape times being associated with more positive charges. Polypeptides enriched in positive charges exhibited slower escape, consistent with attractive electrostatic interactions with the negatively charged tunnel surface. Conversely, chains with more negative charges experienced stronger electrostatic repulsion, leading to faster escape. The neutral polypeptide showed an average escape time about 50% slower than the fastest escape and 99% faster than the slowest, highlighting the critical role of electrostatic interactions in modulating escape dynamics (16). Additionally, ribosome profiling analyses revealed that proteins with slower escape exhibited higher ribosome density at the stop codon compared to faster ejectors (16). While our results show consistency with both all-atom simulations and experimental measurements, the potential for slow escape times demonstrates the importance of enhancing computationally efficient simulations to explore rare events.

**Figure 6.**
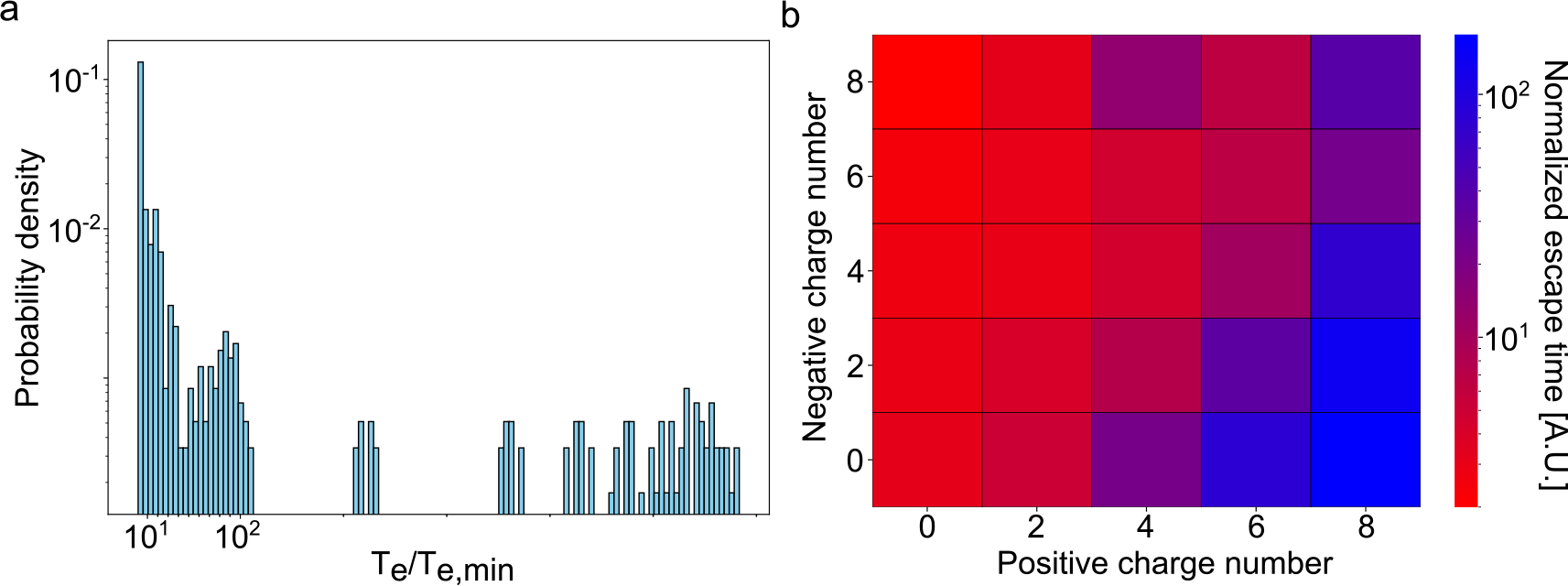
Variations in escape times influenced by charged residues in the C-termini of polypeptides. **(a)** Distribution of normalized escape times for 121 80-mer polypeptide chains simulated using the charged CG ribosome model of 6WD4. Each bar represents the probability density of *T*_*e*_ /*T*_*e*,min_, where *T*_*e*_ is the escape time for each simulation run, and *T*_*e*,min_ is the shortest escape time among all simulations. **(b)** Normalized mean escape times for 25 charge patterns mapped in a heat map. Each cell, except for the neutral protein, represents the average escape time of five charged sequences with randomly distributed charges within the C-terminal 30 residues, normalized by *T*_*e*,min_. Faster and slower escapes are indicated in red and blue, respectively.

## DISCUSSION

In this paper, we present a pipeline for constructing coarse-grained models of the NPET and ribosome surface for MD simulations of nascent polypeptide chains across various contexts and extended timescales. While traditional methods model the NPET interior using probe-based cavity extraction (55, 62–64), our approach quantizes the volume space into a grid to identify empty voxels. This space-partitioning strategy has recently proven effective for generating highly detailed volumetric mesh representations of biomolecules (65) with a highly parallelizable algorithm that leverages octree data structure. Herein, we leverage KDtree structures due to their available Python implementations for voxel querying.

We also note that while our simulations were conducted on unoptimized CPU environments, substantial computational efficiency improvements can be expected through GPU or accelerator packages (38). We employed a simplified CG protein model with Lennard-Jones interactions to illustrate the nascent chain dynamics in our CG environment. While this model is not designed to capture realistic folding, more sophisticated models using established force fields (66) or all-atom representations can explicitly account for side-chain interactions and dynamic behaviors, thereby providing more detailed and accurate representations of translation and folding dynamics.

One significant limitation of our CG model is its inability to capture potentially flexible elements of the NPET and ribosome (such as the lateral stalk (68)) or conformational changes occurring during different steps of translation (69, 70). Interestingly, Kurkcuoglu *et al*. employed elastic network modeling to investigate ribosomal flexibility (71), identifying three distinct tunnel regions with characteristic harmonic motions: the upper region primarily moving parallel to the tunnel axis, the exit region displaying pronounced perpendicular mobility (potentially facilitating nascent peptide chain dispersion), and the constriction site exhibiting rotational motions that may contribute to polypeptide gating mechanisms. Future work could integrate these insights through a hybrid modeling approach where functionally critical flexible regions—particularly at the constriction site—are represented at atomic resolution. This would maintain computational efficiency while incorporating essential ribosomal dynamics, providing a more complete representation of the translation environment and its impact on nascent chain.

## AUTHOR CONTRIBUTIONS

KDD and SS designed the research. AK, SY, ET and WZ implemented the methods. SY carried out all simulations and analyzed the data. SY, AK, KDD and SS wrote the article.

## ACKNOWLEDGMENTS

This research is supported by NSERC Discovery Grants RGPIN-2020-05348 and RGPIN-2024-04666.

## SUPPLEMENTARY MATERIAL

Our SI file contains Supplementary methods and figures, and description of the supplementary videos.

