## Supplementary Information for "Advanced coarse-grained model for fast simulation of nascent polypeptide chain dynamics within the ribosome"

The Supplemental Information contains:

- Supplementary Methods
- Supplementary Figures S1–S4
- Supplementary Videos S1–S4

### 1 Supplementary Methods

#### Location of the PTC and constriction site

In our pipeline, we represent the peptidyl transferase center (PTC) and constriction site as landmarks associated with the nascent polypeptide exit tunnel (NPET), whose coordinates are evaluated as follows:

- **PTC**: We compute the center of mass of LSU rRNA nucleotide, which is closest to the C-terminal residue of the tRNA in the P-Site. The LSU rRNAs and tRNAs of structures **8CCS**, **8HKY**, **8UD8** and **7A5F** are used as templates for this lookup in eukarya, archaea, bacteria and mitochondria respectively.

- **Constriction Site**: We compute the midpoint between the centers of mass of the closest pair of residues in proteins **uL22** and **uL4** in cytosolic structures and **uL22m** and **uL4m** in mitochondrial structures.

##### DBSCAN Parameters

The DBSCAN algorithm clusters datapoints in n-dimensional space based on their density and according to a provided distance metric. The algorithm requires two key parameters: epsilon ($\varepsilon$), which defines the maximum distance between two points for them to be considered neighbors, and *min_samples*, which defines the minimum number of neighbors a point must have to be considered a core point of a dense cluster.

Since our points are simple are simple coordinates of voxels in 3D space we naturally use the euclidian distance metric. We employed a two-pass approach to robustly identify and refine the space representing the NPET. In the first pass, a large neighborhood radius ($\varepsilon = 5.5$ Å) and a high density requirement ($min\_samples = 600$) were used. This combination is effective for identifying the main contiguous void corresponding to the NPET and separating it from smaller, disconnected pockets elsewhere in the ribosome structure. In the second pass, we focused exclusively on this main cluster and used a smaller radius ($\varepsilon = 3$ Å) and a lower density requirement ($min\_samples = 120$). This second step serves to refine the tunnel surface, capturing finer atomic-scale geometric features and removing outlier voxels to produce a smoother and more accurate final mesh.

#### Coarse-grained models of the tunnel and ribosome surface

The NPET and ribosome surface are modeled as purely repulsive walls defined by

$$V_{wall}(r) = \begin{cases} 4\varepsilon\left[\left(\frac{\sigma}{r}\right)^{12} - \left(\frac{\sigma}{r}\right)^{6}\right], & \text{for } r < 3.5 \text{ Å} \\ 0, & \text{for } r \geq 3.5 \text{ Å} \end{cases}$$

where $r$ is the distance from the polypeptide chain to the NPET and ribosome surface measured in angstroms. The parameters of the Lennard-Jones (LJ) potential $\sigma$ and $\varepsilon$ are set to 3.5 Å and 0.066 kcal/mol, respectively, adapted from OPLS-UA force field parameters (1).

Intramolecular interactions between the coarse-grained (CG) beads of the ribosome model are excluded.

The electrostatic interactions between the CG model and the polypeptide chain are simulated using a screened potential from the Debye-Hückel theory,

$$V_{el}(r) = \frac{Cq_iq_j}{\epsilon r} exp(-\kappa r) \quad \text{for } r < 10 \text{ Å}$$

where $r$ represents the distance between the charged beads $i$ and $j$ from the ribosome model and the polypeptide chain, respectively. The dielectric constant $\epsilon$ is set to 7 to account for an implicit solvent system involving ribosomal proteins, rRNAs and water, and the inverse of the Debye length $\kappa$ is set to 0.14 Å$^{-1}$.

#### Coarse-grained model of polypeptide chain

The polypeptide chain is simulated in a bead-spring model in which each amino acid (aa) residue is modeled by a triplet of beads representing the backbone units. For simplicity, all beads is treated as identical as the $C_\alpha$ atom in Glycine. The short-range intermolecular interactions between non-bonded residues are modeled using LJ potential,

$$V_{ij}(r) = \begin{cases} 4\varepsilon\left[\left(\frac{\sigma}{r}\right)^{12} - \left(\frac{\sigma}{r}\right)^{6}\right], & \text{for } r < 12 \text{ Å} \\ 0, & \text{for } r \geq 12 \text{ Å} \end{cases}$$

where the $\sigma$ and $\varepsilon$ are set to 3.5 Å and 0.066 kcal/mol, respectively. $r$ is the distance between the centers of two interacting beads. Intramolecular interactions are modeled with a harmonic bond potential between neighboring beads, defined as

$$V_b = k_b (r_{i,i+1} - r_0)^2 ,$$

where $r_{i,i+1}$ is the distance between the centers of two consecutive beads and $r_0$ is their equilibrium distance. $k_b$ is set to 268 kcal/molÅ$^2$ and $r_0$ is set to 1.529 Å to reflect an effective distance between bonded beads representative a single carbon-carbon bond. A harmonic angle potential is used to model bond angles between consecutive residues, given by

$$V_\theta = k_\theta(\theta - \theta_0)^2,$$

where $k_\theta$ is set to 58.35 kcal/mol and $\theta_0$ was set to 112.7°.

The opls dihedral potential is also used, given by

$$V_\phi = 1/2K_1(1+cos(\phi)) + 1/2K_2(1-cos(2\phi)) + 1/2K_3(1+cos(3\phi)) + 1/2K_4(1-cos(4\phi)),$$

where $K_1$, $K_1$, $K_1$ and $K_1$ are set to 1.3, -0.05, 0.2 and 0 kcal/mol, respectively. The central bead of each triplet is designated as either neutral, positively charged, or negatively charged. The Coulombic pairwise interactions are calculated using a cutoff distance of 12 Å. All parameters are derived from OPLS-UA force field parameters (1).

#### Dynamic simulations of nascent chains

The molecular dynamics (MD) simulations are performed at $T = 310$ K using real units in LAMMPS. The simulation box is set with periodic boundaries. The mass of all CG beads is set to 12 g/mol. The simulation timestep can be set between 1 and 7 fs to maintain stable integration.

##### Post-translational process

To simulate the escape process of polypeptide chains of varying lengths, we align their CG beads along the tunnel centerline, with a distance of 1.529 Å between adjacent beads. The C-terminus (last bead) is positioned at the PTC, while the beads extending beyond the tunnel are placed along the vector from the PTC to the exit port. Initial velocities of the polypeptide chain are set corresponding to the temperature of 310 K, excluding both linear and angular velocities. The system energy is minimized using the steepest descent (sd) algorithm. Following minimization, the system is considered to be equilibrated when the average internal energy remains constant for 50,000 timesteps, requiring at least 100,000 timesteps in total. After equilibration, the C-terminus is released, and the polypeptide chain allows to fold and move freely within the NPET. When the displacement of the C-terminus exceeds the tunnel length ($\sim$ 110 Å), measured using LAMMPS, the simulation continues for an additional 5,000 time steps before termination.

##### Co-translational process

To simulate the protein elongation process, CG beads representing the polypeptide chain are intermittently added at the PTC. The simulation begins with a single bead fixed at the PTC. After 10,000 time steps, this bead is displaced by 0.5 Å along both the x- and z-axes toward the tunnel exit port. A second bead is then introduced at the PTC, and a harmonic bond is created between the two. Following each bead addition, energy minimization is performed using the sd algorithm to prevent overlaps between beads. The newly added bead is fixed at the PTC by excluding its interactions with other particles, while the remainder of the chain is allowed to equilibrate for 10,000 time steps until the internal energy becomes stable. This cycle–bead displacement, new bead insertion, bond (and angle or dihedral for three or more beads, if necessary) formation, energy minimization, and equilibration–is repeated until the desired chain length is achieved. After the last bead is added to the system, the simulation continues for an additional 5,000 time steps before termination. Throughout the simulation, the CG beads of the NPET and ribosome are kept fixed by setting their net forces to zero.

#### 2 Supplementary Figures

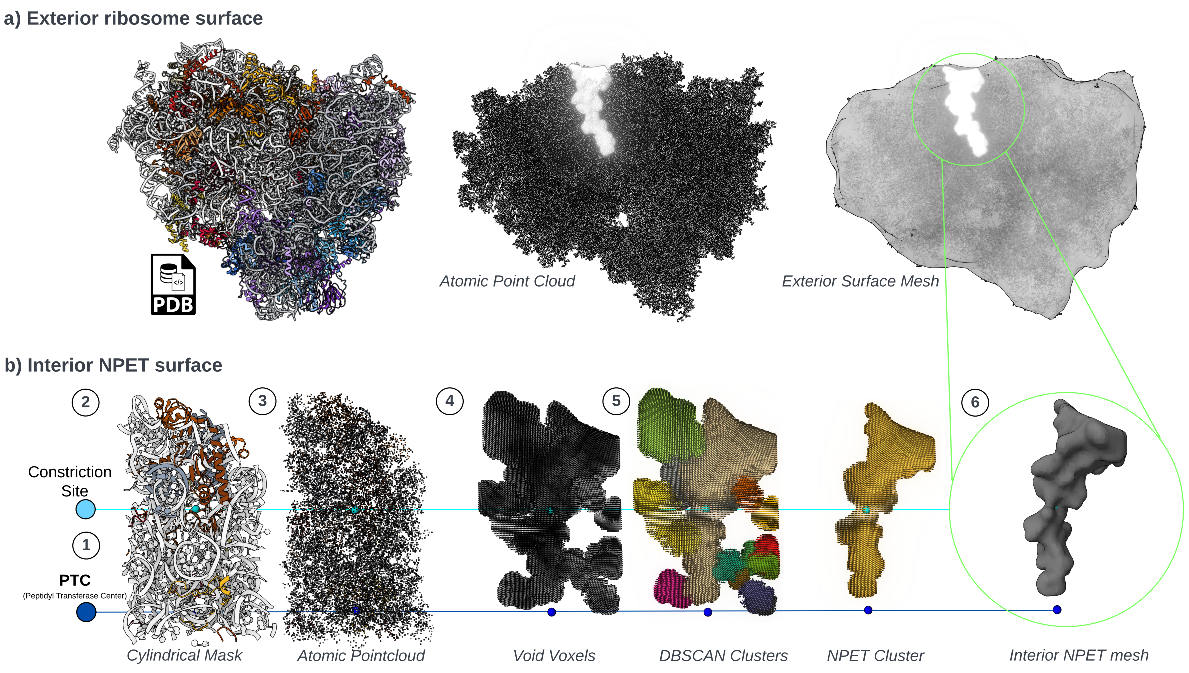


**Figure S1.** Protocol for modeling the **(a)** exterior ribosome surface and **(b)** interior NPET surface, illustrated in six steps (see Methods section).

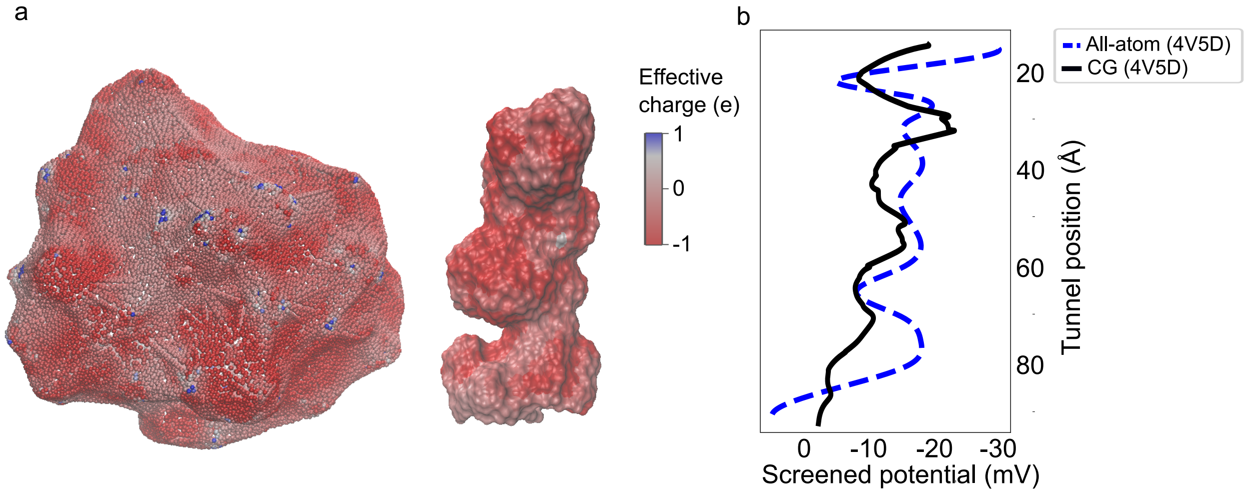


**Figure S2.** Coarse-grained models of the ribosome surface and tunnel with mapping of effective charges. **(a)** Visualization of effective charges mapped onto the ribosome surface and the exit tunnel of 6WD4. Blue and red beads represent positively and negatively charged beads, respectively. The color scale range was adjusted to improve visualization clarity. **(b)** Electrostatic potential along the tunnel centerline in the thermophilic bacterial ribosome 4V5D derived from the CG model (black curve) compared with that from the all-atom simulation model (blue curve) (2).

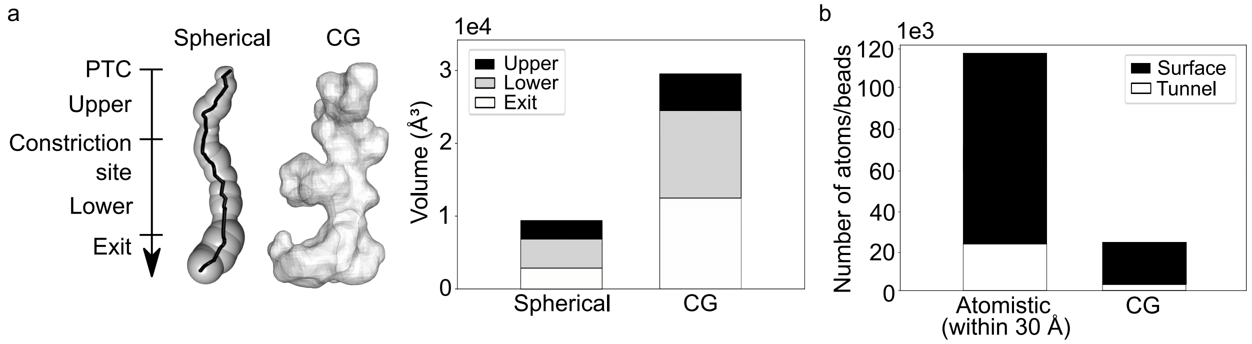


**Figure S3.** Comparisons of the CG tunnel model with the spherical and all-atom model for 6WD4. **(a)** The left panel shows the MOLE-generated spherical tunnel model (left) and the CG model in mesh (right). The tunnel is divided into three regions. The upper, lower and exit region are separated by the constriction site and conserved residues in uL23 or eL39. The right panel compares the volume of the spherical and the CG tunnel models in the upper, lower and exit regions. **(b)** Comparison of the number of beads or atoms in the tunnel (white) and ribosome surface (black) regions for the all-atom and CG ribosome models. The atomistic model includes all atoms within 30 Å of the ribosome surface and the tunnel centerline. In the atomistic model, overlapping atoms between the two regions are excluded in the atom count for the surface region.

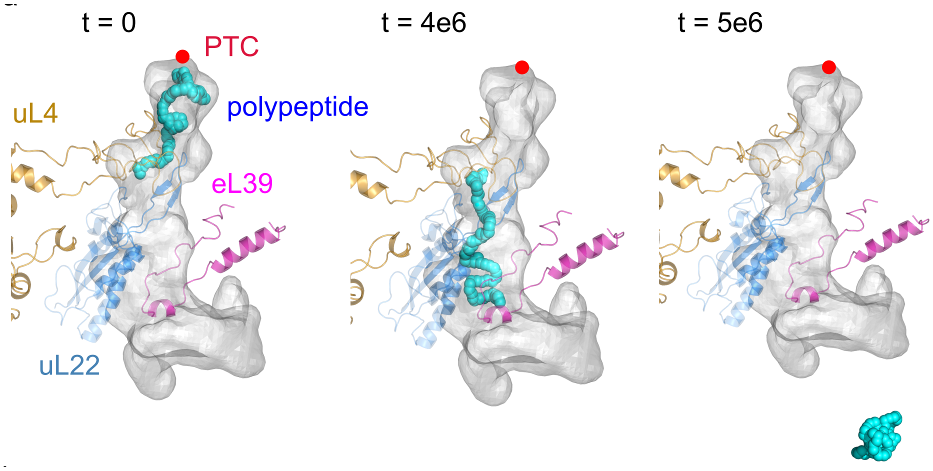


**Figure S4.** Post-translational dynamics of a 20-mer polypeptide, illustrating its release from the PTC to its full exit from the CG tunnel of 4UG0 (left to right). Ribosomal proteins uL4, uL22, and eL39 are highlighted in orange, blue, and magenta, respectively. These proteins are merely used for visualization purposes, and are not explicitly modeled in the simulations. The PTC is highlighted as a red dot. Time t is the simulation timestep in arbitrary unit. The ribosome surface is not shown here.

### 3 Supplementary Videos

The paper includes the following videos.
**Video S1** Visualization of the coarse-grained model of the NPET and ribosome surface of 4UG0.

**Video S2** Comparison of the tunnel volume of the coarse-grained model and the spherical model (generated by MOLE2.5) of 4UG0.

**Video S3** MD simulations of the escape process of the 20-mer polypeptide chain from the exit tunnel of 4UG0.

**Video S4** MD simulations of the elongation process of the 100-mer polypeptide chain within the ribosome of 4UG0.
